# Schizophrenia Polygenic Risk Score and 20-Year Course of Illness in Psychotic Disorders

**DOI:** 10.1101/581579

**Authors:** Katherine G. Jonas, Todd Lencz, Kaiqiao Li, Anil K. Malhotra, Greg Perlman, Laura J. Fochtmann, Evelyn J. Bromet, Roman Kotov

**Author notes:** Address correspondence to: Katherine Jonas, HSC T10-060, Department of Psychiatry Stony Brook University Stony Brook, NY 11794-8101.

## Abstract

Understanding whether and how the schizophrenia polygenic risk score (SZ PRS) predicts course of illness could improve diagnostics and prognostication in psychotic disorders. We tested whether the SZ PRS predicts symptoms, cognition, illness severity, and diagnostic changes over the 20 years following first admission. The Suffolk County Mental Health Project is an inception cohort study of first-admission patients with psychosis. Patients were assessed six times over 20 years, and 249 provided DNA. Geographically- and demographically-matched never psychotic adults were recruited at year 20, and 205 provided DNA. Symptoms were rated using the Schedule for the Assessment of Positive Symptoms and Schedule for the Assessment of Negative Symptoms. Cognition was evaluated with a comprehensive neuropsychological battery. Illness severity and diagnosis were determined by consensus of study psychiatrists. SZ PRS was significantly higher in first-admission than never psychotic groups. Within the psychosis cohort, the SZ PRS predicted more severe negative symptoms (*β* = 0.21), lower GAF (*β* = −0.28), and worse cognition (*β* = −0.35), across the follow-up. The SZ PRS was the strongest predictor of diagnostic shifts from affective to non-affective psychosis over the 20 years (AUC = 0.62). The SZ PRS predicts persistent differences in cognition and negative symptoms. The SZ PRS also predicts who among those who appear to have a mood disorder with psychosis at first admission will ultimately be diagnosed with a schizophrenia spectrum disorder. These findings show potential for the SZ PRS to become a powerful tool for diagnosis and treatment planning.

## Introduction

The SZ PRS agglomerates the weighted effect of many single nucleotide polymorphisms (SNPs) that discriminate schizophrenia cases from healthy controls (1). As the size of the PRS discovery cohort has increased, case-control variance explained has increased from 3% to 18% (1,2). However, there is substantial heterogeneity in both the clinical presentation and illness course of schizophrenia that is difficult to capture in large studies. Carefully phenotyped, longitudinal studies may reveal when and how the SZ PRS’s effects unfold.

Evidence on the association between SZ PRS and clinical presentation has been inconsistent. For example, a study of adults found an association with mood-incongruent psychosis in both schizophrenia and bipolar samples (3), suggesting the PRS is especially sensitive to positive symptoms, while a study of an adolescent cohort found that the SZ PRS predicted negative symptoms and anxiety (4). Still others have found no associations between SZ PRS and symptom dimensions (5,6). Furthermore, although symptom course is a central component of the diagnostic criteria for schizophrenia, the association between the SZ PRS and symptom trajectories in psychotic disorders is unknown. In comparison to links with symptoms, the association of SZ PRS with neurocognitive deficits has been stronger. The SZ PRS has been linked with prodromal motor deficits (7), as well as cognitive decline lower educational attainment in the general population (8,9). However, it is unclear whether SZ PRS predicts either cognition or cognitive decline in people with psychotic disorders (10).

Investigating the longitudinal associations of the SZ PRS is important because diagnosis is often incorrect at first-admission. Due to diagnostic error, cross-sectional studies of first-admission patients may underestimate the magnitude of genetic effects, which gain power as the diagnostic picture clarifies.(11–13). At the time of first onset, the SZ PRS distinguishes patients from controls (14), and predicts treatment response (15). Later on in the course of illness, the SZ PRS can make relatively nuanced distinctions among individuals with schizophrenia, bipolar disorder with psychosis, and bipolar disorder without psychosis (3). However, no one has yet determined whether the SZ PRS can be used in early phases of illness to predict diagnostic course. Investigating the association between the SZ PRS and diagnosis over time may reveal genetic effects that would otherwise be dampened by inaccurate diagnoses, as well as genetic liabilities for specific diagnostic trajectories.

If the SZ PRS is associated with changes in diagnosis, illness severity over time, trajectory of specific symptoms, or cognitive changes, such a pattern would have significant implications for diagnosis, prognosis, and treatment. Given that trajectories of psychotic disorders differ substantially across patients (16,17), the SZ PRS may help to identify patients who are at risk of a worsening course of illness, and who need early and intensive attention to pre-empt deterioration. Alternatively, if the effects of common genetic variants are observable by baseline hospitalization and remain constant across the course of illness, the SZ PRS may shape the prodromal phenotype. The present study addresses these hypotheses by investigating the contributions of the SZ PRS to trajectories in symptom and illness severity, cognition, and diagnosis over 20 years.

## Methods

### Sample

Data were drawn from the Suffolk County Mental Health Project, a longitudinal first-admission study of psychosis. Between 1989 and 1995, individuals with first-admission psychosis were recruited from the 12 inpatient facilities in Suffolk County, New York (response rate 72%). The Stony Brook University Committee on Research Involving Human Subjects and the review boards of participating hospitals approved the protocol annually. Written consent was obtained from all study participants or their parents, for those who were minors at baseline. Eligibility criteria included residence in Suffolk County, age between 15 and 60, ability to speak English, IQ > 70, first admission within the past 6 months, current psychosis, and no apparent medical etiology for psychotic symptoms.

A total of 628 participants met inclusion criteria. Follow-up interviews were conducted at 6 months, 24 months, 48 months, 10 years, and 20 years after baseline. Ninety-two participants died during the follow up period. Of those surviving, 373 were interviewed at the 20 year follow-up (18). Of those 373, DNA was collected from 249 participants as part of the Genomics Psychiatry Cohort collaborative (19).

At the 20 year point, a comparison group of 261 (205 provided DNA) never psychotic adults was recruited using random digit dialing in zip codes where members of the psychosis cohort resided (20). Rate of participation in this group was 67%. The comparison group was sex- and age-matched to the psychosis cohort. Table 1 reports demographics for both the psychosis and never psychotic cohorts.

**Table 1.** *Demographic characteristics*

|  | Psychosis Cohort (N=249) |  | Never Psychotic Adults (N=205) |  |
| --- | --- | --- | --- | --- |
|  | N | % | N | % |
| Gender |  |  |  |  |
| Male | 142 | 57.0 | 118 | 57.6 |
| Female | 107 | 43.0 | 87 | 42.4 |
| Race/Ethnicity |  |  |  |  |
| Caucasian | 208 | 83.6 | 184 | 89.8 |
| African-American | 27 | 10.8 | 9 | 4.4 |
| Other | 14 | 5.6 | 12 | 5.8 |
| Age at 20-year follow up |  |  |  |  |
| 30-40 | 54 | 21.7 | 32 | 15.6 |
| 40-50 | 106 | 42.6 | 80 | 39.0 |
| 50-60 | 63 | 25.3 | 59 | 28.8 |
| 60-70 | 24 | 9.6 | 33 | 16.1 |
| >70 | 2 | 0.8 | 1 | 0.5 |
| Diagnosis |  |  |  |  |
| Schizophrenia | 100 | 40.2 | - | - |
| Schizoaffective Disorder | 34 | 13.7 | - | - |
| Substance-Induced | 13 | 5.2 | - | - |
| Psychosis |  |  |  |  |
| Other Psychoses | 16 | 6.4 | - | - |
| Major Depression | 21 | 8.4 | - | - |
| Bipolar Disorder | 65 | 26.1 | - | - |
| AP Medication | 142 | 57.0 | 4 | 2.0 |
| Illness Severity (GAF) |  |  |  |  |
| <20 | 2 | 0.8 | - | - |
| 21-40 | 115 | 46.2 | - | - |
| 41-60 | 70 | 28.1 | 31 | 15.1 |
| 61-80 | 46 | 18.5 | 112 | 54.6 |
| >80 | 16 | 6.4 | 62 | 30.3 |

*Note.* GAF = Global Assessment of Functioning.

For genetic assays, DNA was extracted from peripheral lymphocytes and genotyped using the Illumina PsychArray-8 platform containing 571,054 markers. Standard quality control procedures were performed to exclude SNPs with minor allele frequency (MAF) <1%, genotyping failure >5%, Hardy-Weinberg equilibrium p<10^−6^, mismatch between recorded and genotyped sex, as well as related individuals (π̂>.20, in which case the relative with the lower call rate was dropped). Mean call rate was 99.8%. SNP imputation was conducted with IMPUTE2 (21), against the full 1000 Genomes phase 3 reference panel (22). The imputed SNPs underwent another round of quality control and SNPs with missing data >5% and imputation information score <0.8 were excluded, yielding 6.87M high quality biallelic SNPs. Genomic data analysis was performed using the SVS software, version 8.7.0 (23).

The PRS was calculated for each participant in the sample as the weighted sum of the risk allele they carried, based on the summary statistics (effect alleles and odds ratios) derived from the clumped PGC-2 GWAS results, which consists of 102,636 SNPs (2). The clumped PGC GWAS summary statistics file was downloaded from the LD Hub at the Broad Institute (http://ldsc.broadinstitute.org/ldhub/). The clumping parameters are as following: a SNP will be clumped to a more significant SNP with LD (r^2^ ≥ 0.10) within a 500kb window. PRS calculation was carried out in the PRSice software (24). The SZ PRS was calculated at several p-value thresholds (p ≤ 5×10^−8^, 0.001, 0.01, 0.05, 0.10, 0.20, and 0.50). Mean PRS scores at these thresholds for each diagnostic group are reported in Supplemental Table 1. The results presented here utilize PRS scores based on SNPs with a p-value < 0.01, as this threshold provided the greatest separation between the diagnostic and never-psychotic groups. However, analyses scores based on other thresholds yielded similar results (available upon request).

### Measures

#### Diagnosis

Research diagnoses were made by the consensus of study psychiatrists at baseline and again at year 20 using all available longitudinal information, including results of the SCID (25), interviews with participants’ significant others, medical records, and observations and behavioral ratings by master-level interviewers (the diagnostic process is described in Bromet, 11).

#### Symptoms

Symptom domains were rated using the Scale for the Assessment of Positive Symptoms (SAPS; 26), and the Scale for the Assessment of Negative Symptoms (SANS; 27). Masters-level mental health professionals made ratings of symptoms based on their interview of the participant, interviews with significant others, and medical records. SAPS and SANS ratings were used to score four factor-analytically derived subscales: reality distortion (delusions/hallucinations; Cronbach’s α=0.85) and disorganization (α=0.77) from the SAPS; avolition (α=0.87) and inexpressivity (α=0.90) from the SANS (28). Sample sizes at each time point were as follows: 6 months (N=217); 24 months (N=209); 48 months (N=192); 10 years (N=234); and 20 years (N=246). Depression in the past month was rated via the Structured Clinical Interview for DSM-III-R at baseline (25) and DSM-IV thereafter (29), administered without skip-outs and scored as a sum of 9 symptom ratings. The Global Assessment of Functioning (GAF) was used to rate overall illness severity (symptoms plus functional impairment) by the consensus of study psychiatrists using all available information.

#### Cognition

The neuropsychological battery at 24-month and 20-year follow-up included Verbal Paired Associates and Visual Reconstruction (WMS-R, immediate and delay trials; 30), Symbol-Digit Modalities (WAIS-III; 31), Trails A and B (32), the Controlled Oral Word Association Test (33), Vocabulary (WRAT-3; 34), and the Stroop Test (35). Sample sizes for the first-admission cohort were N=201 at 24 months and N=224 at 20 years.

### Analyses

#### Attrition

Supplemental Table 2 describes the sample sizes available for each outcome measure at each time point, as well as Cohen’s *d* comparing outcome measures between those who did and did not provide DNA. Those who provided DNA were slightly younger (Cohen’s *d=*-0.18, p < 0.05), more likely to be prescribed antipsychotic medications at 24 months (Cohen’s *d=0.14*, p < 0.05), 48 months (Cohen’s *d*=0.17, p < 0.05), and 20 years (Cohen’s *d=*0.17, p < 0.05), had higher ratings on SANS avolition at baseline (Cohen’s *d=*0.18; p < 0.05), and lower ratings at 48 months (*d=*-0.23; p < 0.05), and had better scores on the COWAT (*d*=0.60, p < 0.05). All analyses used full information maximum likelihood estimation methods, which use all data, including partial cases, to arrive at unbiased parameter estimates.

**Table 2.** *Symptom trajectories over 20 years*

|  | Mean Effect of<br>SZ PRS |  | SZ PRS on<br>Course Part 1 |  | SZ PRS on<br>Course Part 2 |  |
| --- | --- | --- | --- | --- | --- | --- |
| | $\beta$ | SE | $\beta$ | SE | $\beta$ | SE |
| Hallucinations/Delusions | 0.04 | 0.08 | 0.01 | 0.06 | NA | NA |
| Disorganization | 0.09 | 0.09 | 0.02 | 0.07 | 0.03 | 0.08 |
| Avolition | 0.21* | 0.09 | -0.03 | 0.05 | -0.01 | 0.05 |
| Inexpressivity | 0.14 | 0.09 | -0.07 | 0.05 | -0.05 | 0.06 |
| Depression | 0.12 | 0.08 | -0.10 | 0.05 | NA | NA |
| Illness severity | -0.28** | 0.09 | -0.03 | 0.05 | 0.04 | 0.04 |

*Note.* All effects are standardized coefficients ( $\beta$ s). The mean effect of the SZ PRS indicates an effect that is constant across all time points. SZ PRS on course is the effect of the SZ PRS on change in symptoms over time. For Reality Distortion and Depression, symptom trajectories are linear, so there is only one effect on slope. Trajectories have inflection points for Disorganization (13.5 years after baseline), Avolition (4.5 years), Inexpressivity (2.5 years), and Illness severity (4.5 years), so effect of PRS is also modeled on the second slope for these domains. \* $p < 0.05$ ; \*\* $p < 0.01$ .

#### Baseline associations

The baseline time point was qualitatively different from follow-up time points, so baseline symptoms and trajectories from 6 months to 20 years were analyzed separately. Associations of the SZ PRS with baseline symptoms and illness severity were tested via linear regression adjusted for age.

#### Trajectories

Multilevel spline regression models were used to estimate trajectories of symptoms and illness severity. To allow for non-linear trajectories, we identified the point at which the average trajectory changed direction. The placement of the change point was determined by alternatively placing it at each 1-year interval from baseline to 20-year follow-up, and comparing the fit of these competing models via the Bayesian Information Criterion (BIC).

#### Cognitive change

Parallel analysis was used to determine how many factors were reflected in the set of cognitive tests administered at the 24-month and 20-year follow-ups. In both cases, parallel analysis indicated a single cognitive factor. One-factor confirmatory measurement models were fit to the cognitive tests at each time point. In both the 24-month and 20-year models, all test loadings on the general factor were greater than 0.3 and statistically significant. Residual covariance terms were included between subscales of tests with more than one subscale. Model fit was excellent at both 24 months (CFI=0.94; RMSEA=0.04) and 20 years (CFI=1.00; RMSEA=0.01). The latent cognition factor was regressed on the PRS and age.

#### Diagnostic shifts

Baseline and 20 year diagnoses were dichotomized into non-affective psychosis and affective psychosis as described in Kotov (36), which was consistent with similarity of PRS scores among specific diagnoses. Supplemental Table 3 reports the SZ PRS scores for diagnostic groups at baseline and 20-year follow-up relative to never psychotic adults. The affective psychosis (AP) category included psychotic bipolar disorder and psychotic major depression. The non-affective psychosis (NAP) category included schizophrenia, schizoaffective disorder, substance-induced psychosis, and other psychoses. Contrasts between diagnostic groups and shift groups were calculated by regressing PRS scores on group status. Statistical significance was determined by non-parametric permutation and bootstrap tests.

Predictive modeling of shifts from AP to NAP between baseline and 20 years was performed by regressing diagnostic shift groups on statistically significant clinical predictors from Table 3 of Bromet (11). Jackknife resampling and leave-one-out cross-validation were used to calculate the stability of model estimates and prediction error, respectively.

#### Population stratification

In order to control for population stratification due to ancestry, all analyses were covaried on the first ten principal components of genetic covariance (37). In addition, we completed sensitivity analyses on the subsample of Caucasian participants. Because the PGC-2 SZ PRS was calibrated in a largely Caucasian sample, it is less accurate in non-Caucasian samples (38). We therefore replicated these analyses in the subset of Caucasian participants (N = 392). These results are reported in Supplemental Material.

#### Multiple comparisons

To limit the number of Type I errors, we employed Benjamini and Hochberg’s procedure for controlling the false-discovery rate at q = 0.10 (39). Among the 46 contrasts completed in these analyses, all reported p-values remain significant after FDR, with one exception noted in Footnote 3.

## Results

### Baseline Associations

The SZ PRS was not associated with severity of hallucinations and delusions, disorganization, or inexpressivity at first admission. There was an effect of the PRS on avolition (β=0.33, SE=0.12, p < 0.01). The PRS was also not associated with depression or global severity.

### Symptom Trajectories

Table 2 reports the associations between the SZ PRS and the symptom domains of schizophrenia, as well as depression and global illness severity. The SZ PRS was not associated with hallucinations/delusions, disorganization, inexpressivity, or depression. Rather, the SZ PRS was associated with stable differences in avolition (β=0.21, SE=0.09, p < 0.05) and illness severity (GAF; β=-0.28, SE=0.09, p < 0.01). The SZ PRS did not predict changes in symptoms over time, but stable differences across all time points. Figure 1 depicts the mean trajectory of illness severity and avolition for those with high versus low PRSs.

**Figure 1.**
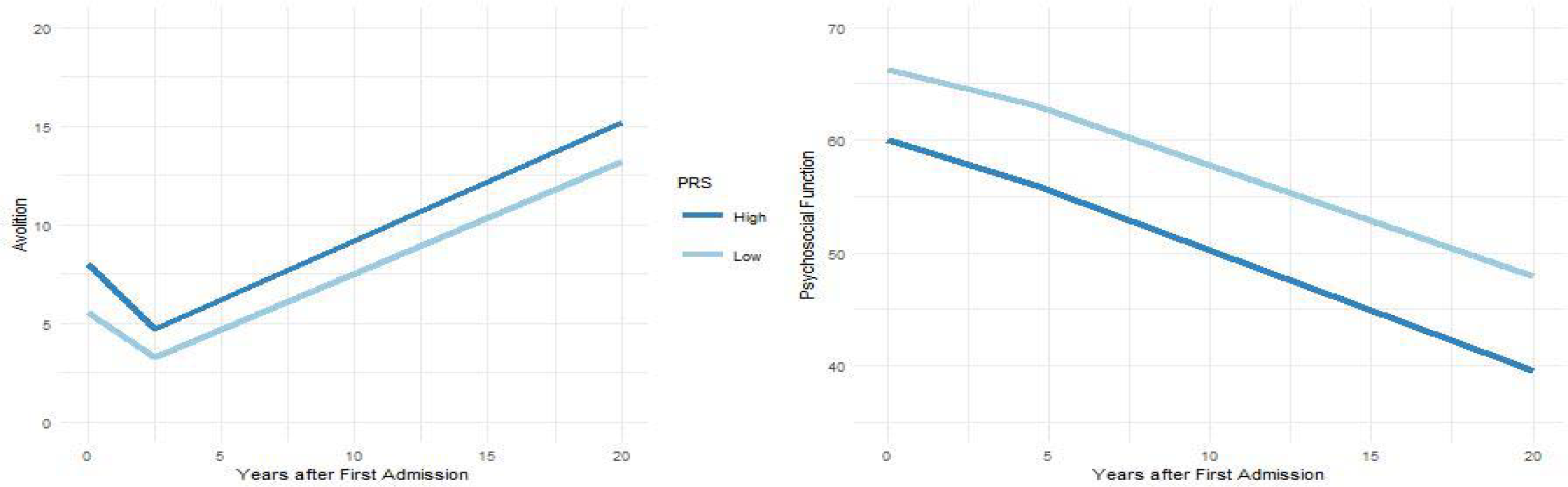
Effect of SZ PRS on course of illness. *Note.* The course of negative symptoms (SANS avolition, left) and illness severity (GAF, right) over the twenty years following first hospitalization. Light and dark blue lines depict the course of illness for those with low and high SZ PRS scores, respectively, as defined by a median split.

### Cognitive Outcomes

SZ PRS was associated with poorer cognition at 24 months (β=-0.29, SE=0.13, p < 0.05) and at 20 years (β=-0.35, SE=0.11, p < 0.01), though not with change between the two time points. Given the correlation between negative symptoms and cognition (r=-0.32 at 24 months, and r=-0.44 at 20 years), we adjusted the models for avolition, but this did not change the pattern of effects at 20 years.^1^

### Diagnostic Shifts

We next tested whether the SZ PRS predicted shifts between affective psychosis (AP) and non-affective psychosis (NAP). Those in the NAP group at both baseline and 20 years had larger SZ PRS scores than those who were in the AP group at both times points (*d*=0.27, p < 0.05). Notably, those whose baseline AP diagnosis was changed to NAP by the 20-year follow-up had larger SZ PRS scores compared to participants whose diagnosis was AP at both time points (*d*=0.45, p < 0.05; equivalent to AUC=0.62), and were not distinguishable from those who were in the NAP group at both time points (*d*=0.14, p=0.55).^2^ Figure 2 depicts the distribution of SZ PRS scores for these groups. At high levels of the SZ PRS, liability for NAP (either stable or shifted into NAP) versus stable AP was pronounced. At the highest decile of scores, the SZ PRS detects those who will transition into NAP group with 83% accuracy (AUC=0.83; 5 of 11 individuals who transitioned).

**Figure 2.**
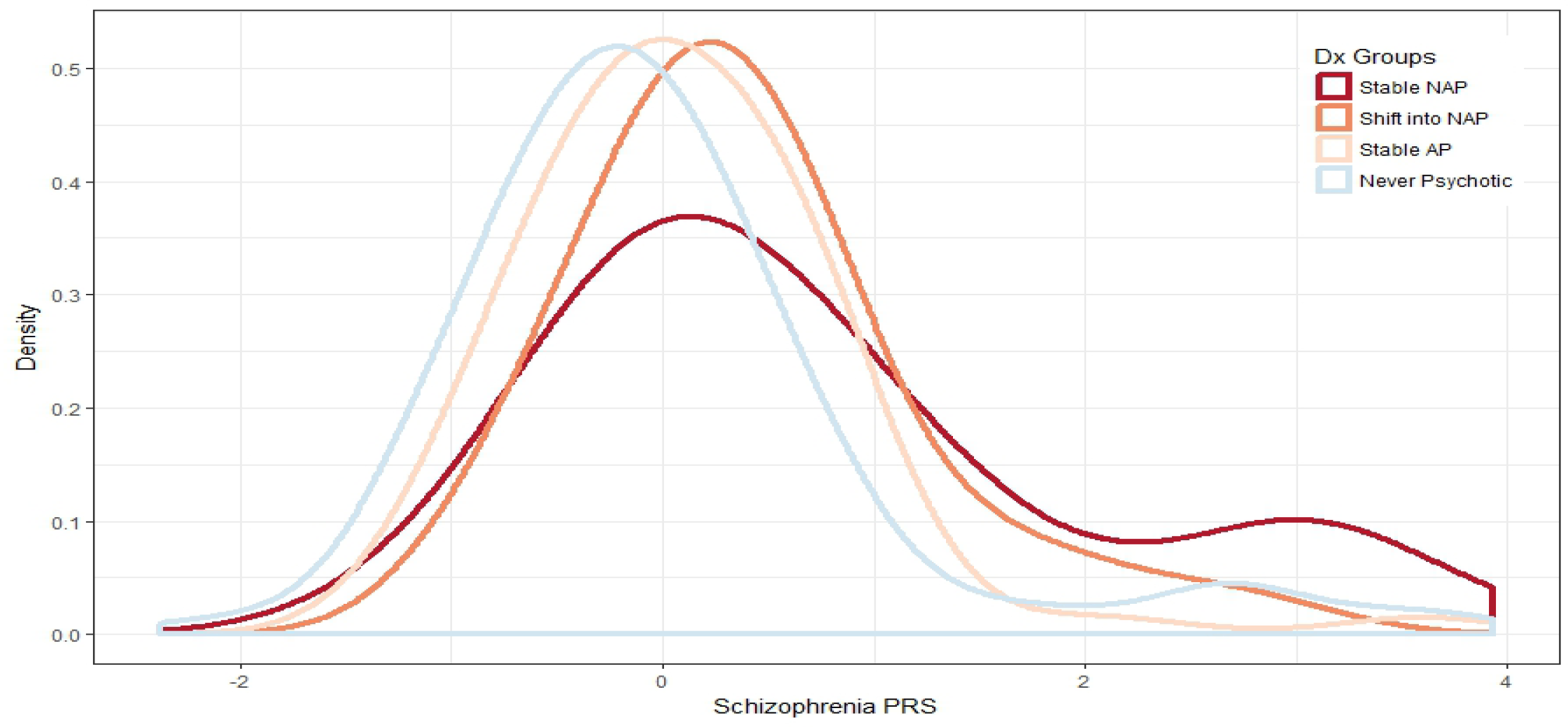
Distribution of SZ PRS scores in diagnostic shift groups. *Note.* AP is affective psychosis; NAP is non-affective psychosis. Density represents the probability of observing a given PRS score within a given diagnostic shift group. All SZ PRS scores are standardized relative to never psychotic adults.

To compare the performance of the SZ PRS to clinical predictors, we added significant predictors from Bromet (11)—baseline GAF, negative, psychotic, disorganized, depressive, and manic symptoms; antipsychotic and mood stabilizers prescriptions—to the regression described above. In the resulting model, only treatment with mood stabilizers (AUC=0.63) and the SZ PRS (AUC=0.61) predicted a shift from AP to NAP (total AUC=0.70; leave-one-out prediction error=0.33).^3^

## Discussion

As the size of the calibration sample for the SZ PRS has increased, so has the power to distinguish cases and controls (14,40). However, it has remained unclear how the SZ PRS influences psychotic disorders over time. Taken together, these analyses show that the SZ PRS is not only a robust predictor of diagnosis, but also of illness severity and cognitive deficits over time. We found that the SZ PRS predicts a course of persistent negative symptoms. Importantly, among patients initially diagnosed with a psychotic affective disorder, a higher SZ PRS predicted whose diagnosis would change to non-affective psychosis by the 20-year follow-up. This effect was modest (AUC*=*0.61), but was stronger in the highest decile (AUC=0.83). Indeed, clinical predictors of this shift improved on the SZ PRS only modestly, increasing AUC from 0.62 to 0.70.

We tested the association of the SZ PRS with overall illness severity and the major symptom domains (hallucinations/delusions, disorganization, avolition, inexpressivity, and depression). Our results suggest that SZ PRS is more closely linked to avolition than other symptom domains. Furthermore, the SZ PRS establishes a set degree of severity relative to others with the same illness, rather than different illness trajectories. We found similar associations between the SZ PRS and cognition. The SZ PRS predisposes people with psychotic disorders to consistently worse cognition—specifically, a one standard deviation increase in PRS would predict a 5-point lower IQ—rather than to cognitive decline after illness onset. This is consistent with cognitive deficits observed in high risk cohorts who have not yet had a psychotic episode (41), as well as the effect of the cognitive PRS in schizophrenia cohorts (42).

Furthermore, the stability of the SZ PRS’ effects after first admission suggests genetic liabilities for schizophrenia take their toll in the prodromal phase, or before. Taken together, the findings support a neurodevelopmental continuum model of psychosis, such that increased genetic burden predicts cognitive deficits and negative symptoms that emerge prior to eventual diagnosis of non-affective psychosis (43). Attempts to identify mechanisms through which the SZ PRS exerts its effects should focus on prodromal samples.

Allardyce and colleagues (3) found the SZ PRS discriminated psychotic bipolar disorder from schizophrenia. Here, we extend that finding, showing that the SZ PRS discriminates more broadly between affective and non-affective psychosis. These findings also parallel those of Vassos and colleagues (14), who show the SZ PRS to be elevated in schizophrenia as compared to other psychotic disorders at first episode. Here, we show that not only is the SZ PRS elevated in schizophrenia in comparison to affective psychosis, but also that an elevated SZ PRS predicts an eventual diagnosis of non-affective psychosis. Taken together, these results suggest the core pathology of schizophrenia—cognitive deficits and negative symptoms (18,44)—were present at first admission but obscured by comorbid mood symptoms. In the months and years following first admission, affective and positive symptoms may subside, while negative and cognitive symptoms persist, resulting in a change of diagnosis to non-affective psychosis.

For common disorders such as coronary artery disease, type 2 diabetes, and breast cancer, PRS scores are already showing potential clinical utility (45). In psychiatric disorders, the accuracy and therefore utility of PRS scores has been limited by low base rates. However, within samples of psychotic individuals, and especially at the extreme tails of the distribution of scores, the SZ PRS may be clinically useful for predicting illness course. These findings show promise for personalized medicine for psychiatric disorders. Furthermore, as the size of the Psychiatric Genomic Consortium sample grows, so will the statistical power of the SZ PRS (40). The SZ PRS may identify a subset of patients with poor prognoses even when the initial diagnostic picture is unclear. Those with especially high SZ PRS scores at first admission are at risk of worse outcomes and may benefit from more intensive follow-up.

### Strengths and Limitations

The design of this study is unique in that it is the only study that has linked course of illness to the SZ PRS. As such, it was uniquely positioned to speak to the correlates of genetic liabilities for schizophrenia as they unfold over time. Still, our analyses are limited in three notable ways. First, of the 628 cases who met inclusion criteria only 249 consented to genetic assays at the 20-year follow-up. An analysis of missing data shows few consistent patterns. Those who provided DNA were more likely to be prescribed antipsychotics across the follow-up period, but this trend is difficult to interpret. The prescription may reflect greater symptom severity, or symptoms may be less severe due to being treated, but neither of these trends were reflected in measures of symptom severity. Second, the sample size is modest (249 subjects with psychotic disorders and 205 who were never psychotic). Repeated clinical assessments over time increased the precision of the estimates, but we do not know whether a larger sample would detect effects that were not found here. Finally, predictive modeling always requires cross-validation in a new sample. While our in-sample analyses suggest the effect of the SZ PRS is robust, replication is needed. Importantly, the primarily Caucasian ancestry of the Psychiatric Genomic Consortium reduces the predictive power of polygenic risk scores in other ancestry groups (38). While the SZ PRS is useful in this sample, which is 85% Caucasian, it will be substantially less so in non-Caucasian individuals. As the PGC develops PRS scores in other ancestry groups, it will be important to determine whether a similar pattern of results holds.

## Conclusions

The SZ PRS predisposes individuals to consistently worse course of illness severity, avolition, and cognitive deficits. Among those patients with a mix of affective and non-affective symptoms at baseline, high SZ PRS scores predicted those whose diagnosis would shift to non-affective psychosis in the 20 years following first-admission. Together, these findings suggest the SZ PRS is an indicator of the core clinical correlates of schizophrenia. They also highlight the potential of the SZ PRS as a diagnostic aid and prognostic indicator for patients with especially high scores, who may benefit from early and consistent monitoring and care.

## Acknowledgements

This research was supported by National Institutes of Health (MH44801 to EB, MH094398 to RK, MH117646 to TL, MH085548 subcontract from Carlos Pato to EB, and MH085542 subcontract from Carlos Pato to EB), and a NARSAD Young Investigator Grant to RK. The authors gratefully acknowledge the support of the participants and mental health community of Suffolk County for contributing their time and energy to this project. They are also indebted to the study coordinators for their dedicated efforts, the interviewers for their careful assessments, and the psychiatrists who derived the consensus diagnoses.

A subset of these analyses were presented at the 30th annual meeting of the Society for Research in Psychopathology, Denver, CO, September 14-17th, 2017.

## Sensitivity Analyses

Results were the same in the subsample of Caucasian participants (N = 392), with the exception of three effects that were no longer significant: One, the association between SZ PRS and avolition at baseline (β=0.13, p = 0.08); Two, contrasts between diagnostic shift groups were no longer significant (stable AP versus stable NAP *d* = 0.27, p = 0.18; stable AP versus shift into NAP *d =* 0.36, p = 0.10). This may be attributable to the 18% reduction in sample size of the NAP group when non-Caucasian participants were removed.

---

**Table S1.**
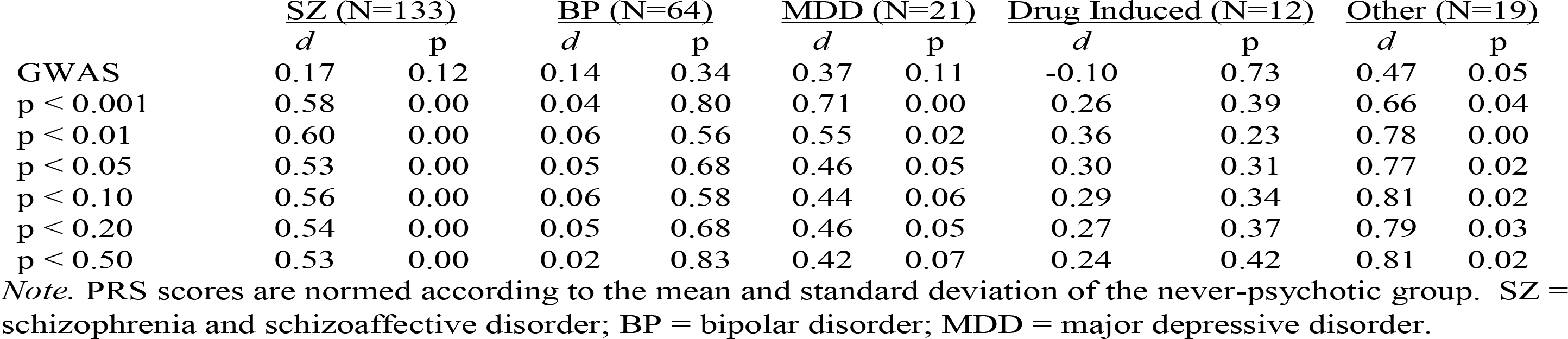
*Effect of PRS thresholds on diagnostic group means*

**Table S2.** *Analysis of patterns of missing data among cases*

|  | BL |  |  | 6 M |  |  | 24 M |  |  | 48 M |  |  | 10 Y |  |  | 20 Y |  |  |
| --- | --- | --- | --- | --- | --- | --- | --- | --- | --- | --- | --- | --- | --- | --- | --- | --- | --- | --- |
|  | N | % | *d* | N | % | *d* | N | % | *d* | N | % | *d* | N | % | *d* | N | % | *d* |
| Demographics |  |  |  |  |  |  |  |  |  |  |  |  |  |  |  |  |  |  |
| Gender | 249 | 39.6% | 0.02 | - | - | - | - | - | - | - | - | - | - | - | - | - | - | - |
| Ethnicity | 249 | 39.6% | 0.10<sup>a</sup> | - | - | - | - | - | - | - | - | - | - | - | - | - | - | - |
| Age | 249 | 39.6% | -0.18* | - | - | - | - | - | - | - | - | - | - | - | - | 249 | 64.7% | -0.11 |
| Diagnosis |  |  |  |  |  |  |  |  |  |  |  |  |  |  |  |  |  |  |
| NAP | 152 | 39.1% | 0.01 | - | - | - | - | - | - | - | - | - | 154 | 38.3% | 0.04 | - | - | - |
| AP | 97 | 40.6% | 0.01 | - | - | - | - | - | - | - | - | - | 95 | 42.0% | 0.04 | - | - | - |
| AP Medication | 249 | 39.6% | .021 | 249 | 39.9% | .034 | 247 | 42.1% | 0.14* | 249 | 40.30% | 0.17* | 234 | 53.9% | .095 | 240 | 76.7% | 0.17* |
| Symptoms |  |  |  |  |  |  |  |  |  |  |  |  |  |  |  |  |  |  |
| Reality Distortion | 249 | 39.6% | -0.09 | 217 | 46.3% | -0.12 | 209 | 49.6% | -0.01 | 192 | 53.8% | -0.11 | 234 | 53.9% | -0.17 | 246 | 74.1% | 0.12 |
| Disorganization | 249 | 39.6% | 0.13 | 217 | 46.3% | 0.04 | 209 | 49.6% | -0.07 | 192 | 53.9% | -0.07 | 222 | 58.4% | 0.03 | 246 | 75.5% | 0.12 |
| Inexpressivity | 248 | 39.6% | 0.14 | 217 | 46.3% | -0.06 | 209 | 49.6% | -0.01 | 191 | 53.8% | -0.21 | 219 | 61.3% | -0.17 | 239 | 91.2% | -0.32 |
| Avolition | 248 | 39.6% | 0.18* | 217 | 46.3% | 0.02 | 209 | 49.6% | -0.10 | 192 | 53.9% | -0.23* | 234 | 53.8% | -0.16 | 244 | 74.2% | 0.10 |
| Depression | 249 | 39.6% | 0.04 | 217 | 46.4% | 0.03 | 209 | 49.9% | 0.06 | 192 | 53.5% | 0.01 | 231 | 55.4% | 0.05 | 233 | 74.2% | 0.15 |
| Illness Severity | 249 | 39.7% | -0.12 | 221 | 45.1% | 0.02 | 222 | 45.9% | 0.02 | 224 | 48.2% | 0.09 | 238 | 52.9% | 0.10 | 239 | 66.0% | -0.02 |
| Cognition |  |  |  |  |  |  |  |  |  |  |  |  |  |  |  |  |  |  |
| VP1 | - | - | - | - | - | - | 191 | 51.9% | 0.00 | - | - | - | - | - | - | 224 | 94.9% | 0.10 |
| VP2 | - | - | - | - | - | - | 188 | 51.8% | 0.12 | - | - | - | - | - | - | 221 | 95.3% | -0.12 |
| VR1 | - | - | - | - | - | - | 200 | 51.7% | 0.02 | - | - | - | - | - | - | 220 | 94.8% | 0.11 |
| VR2 | - | - | - | - | - | - | 198 | 51.6% | -0.09 | - | - | - | - | - | - | 216 | 94.7% | 0.18 |
| SD | - | - | - | - | - | - | 193 | 50.9% | 0.06 | - | - | - | - | - | - | 223 | 94.9% | 0.43 |
| TMT A | - | - | - | - | - | - | 198 | 51.4% | 0.04 | - | - | - | - | - | - | 215 | 94.7% | -0.13 |
| TMT B | - | - | - | - | - | - | 191 | 51.2% | 0.02 | - | - | - | - | - | - | 212 | 94.6% | -0.34 |
| COWAT | - | - | - | - | - | - | 194 | 51.6% | 0.01 | - | - | - | - | - | - | 219 | 94.8% | 0.60* |
| Vocab. | - | - | - | - | - | - | 201 | 51.3% | -0.02 | - | - | - | - | - | - | 217 | 94.3% | 0.15 |
| Stroop | - | - | - | - | - | - | 183 | 50.8% | 0.05 | - | - | - | - | - | - | 185 | 93.9% | 0.58 |

*Note.* Contrasts are between those who provided DNA (coded as 1) versus those who did not. BL = Baseline; NAP = Non-Affective Psychosis; AP = Affective Psychosis; AP Medication = Antipsychotic Medication; VP1 = Verbal Paired Associates, Immediate Recall; VP2 = Verbal Paired Associates, Delayed Recall; VR1 = Visual Reconstruction,
Immediate Recall; VR2 = Visual Reconstruction, Delayed Recall; SD = Symbol-Digit Modalities; TMT A = Trail Making Test, part A; TMT B = Trail-Making Test, part B; COWAT = Controlled Oral Word Association Task; Vocab. = Vocabulary; $d$ = Cohen's $d$ ; <sup>a</sup> Effect size is Cramer's $V$ , a measure of association for nominal variables; \* $p < 0.05$ ;

**Table S3.** *PRS scores by diagnostic group*

| Diagnosis | Baseline |  | 20 years |  |
| --- | --- | --- | --- | --- |
|  | N | Cohen's *d* | N | Cohen's *d* |
| Schizophrenia/Schizoaffective | 73 | 0.56** | 133 | 0.68** |
| Other Psychoses | 67 | 0.65** | 19 | 0.50* |
| Substance-Induced Psychosis | 12 | 0.86 | 12 | 0.28 |
| Bipolar Disorder | 62 | 0.47** | 64 | 0.36* |
| Major Depression | 35 | 0.45* | 21 | 0.60* |

*Note.* Cohen's *d* is calculated relative to never-psychotic adults. \* $p < 0.05$ ; \*\* $p < 0.01$ .

## Footnotes

1 The model of cognition at 24 months failed to converge when adjusted for avolition.

2 These patterns remained the same when participants with substance-induced psychosis were removed from the analysis (stable NAP versus stable AP *d*=0.30, p<0.05; stable NAP versus shift into NAP *d*=0.07, p=0.82), with the exception of the contrast between stable AP and shift into NAP groups (*d*=0.38, p=0.06).

3 In this prediction analysis, the effect of mood stabilizers, but not the SZ PRS, survived control of the FDR (SZ PRS p-value = 0.04, critical value = 0.03).

